# CAMK2-NR4A1 signaling initiates metabolic substrate switching to induce heart failure with reduced ejection fraction

**DOI:** 10.1101/2025.10.16.682731

**Authors:** Alireza Saadatmand, Mark Pepin, Zihao Chen, Joshua Hartmann, Qiang Sun, Matthias Dewenter, Sumra Nazir, Harikrishnareddy Paluvai, Marco Hagenmüller, Uwe Haberkorn, Lisa Schlicker, Roberto Carlos Frias-Soler, Jens Tyedmers, Christoph Maack, Almut Schulze, Johannes Backs

**Affiliations:** Heidelberg University, Medical Faculty Heidelberg, Institute of Experimental Cardiology, 69120 Heidelberg, Germany; Heidelberg University Hospital, Department of Internal Medicine VIII, 69120 Heidelberg, Germany; German Center for Cardiovascular Research (DZHK), Partner Site Heidelberg/Mannheim, 69120 Heidelberg, Germany; Brigham and Women’s Hospital, Harvard Medical School, Boston, USA; Heidelberg University, Medical Faculty Heidelberg, Department of Nuclear Medicine, 69120 Heidelberg, Germany; German Cancer Research Center (DKFZ), Division of Tumor Metabolism and Microenvironment, 69120 Heidelberg, Germany; Department of Translational Research, Comprehensive Heart Failure Center (CHFC), University Clinic Würzburg, Würzburg, Germany; Molecular Medicine Partnership Unit, Heidelberg University, 69120 Heidelberg, Germany; Helmholtz Institute for Translational AngioCardioScience (HI-TAC) – a branch of the MDC at Heidelberg University, 69120 Heidelberg, Germany

## Abstract

Heart failure with reduced ejection fraction (HFrEF) is marked by a shift in cardiac energy metabolism from fatty acid oxidation to glucose utilization. This “fuel switch” promotes accumulation of glucose byproducts that modify calcium-handling proteins and impair cardiac function, yet the initiating signals remain unclear. We identify Ca^2+^/calmodulin-dependent protein kinase II (CAMK2) as an upstream regulator that triggers pathological substrate switching leading to cardiac systolic dysfunction. Dynamic [^18^F]FDG-PET imaging showed a six-fold increase in myocardial glucose uptake after pressure overload in control mice, but not in cardiomyocyte-specific *Camk2d/Camk2g* double knockouts (cDKO), even before functional decline. cDKO hearts retained lipid reserves, indicating preserved fatty acid metabolism. Transcriptomics revealed strong CAMK2-dependent induction of Nr4a1 and early repression of genes for fatty acid uptake and β-oxidation preceding upregulation of genes for glucose utilization. Cardiomyocyte-specific *Nr4a1* knockout mice closely mimicked the metabolic protection seen in cDKO, while NR4A1 overexpression in human iPSC-derived cardiomyocytes suppressed fatty acid metabolism. NR4A1 directly bound and repressed the FATP1 (Slc27a1) promoter, thereby secondarily enhancing glucose utilization. Together, these findings define a CAMK2–NR4A1 signaling axis that drives lipid depletion and metabolic remodeling, establishing it as a causal mechanism linking energy substrate switching to HFrEF.

## Introduction

The healthy heart is a metabolically flexible organ capable of utilizing multiple energy substrates including fatty acids, carbohydrates, ketones and amino acids to maintain its high energetic demands [1, 2, 3]. The relative proportions of these individual substrates in ATP production can dramatically change under physiologic scenarios. For instance, in fetal life, the heart obtains most of its energy from carbohydrates [4]. Soon after birth, however, this largely changes to β-oxidation of fatty acids, which is maintained in the healthy adult heart [5]. In heart failure with reduced ejection fraction (HFrEF), cardiac function is reduced, which is accompanied by profound metabolic rewiring, including disturbed energy metabolism and impaired metabolic flexibility, characterized by a switch from fatty acid metabolism back towards a more fetal-like metabolic pattern with preferential use of glucose as main fuel source [6-8]. While it is generally accepted that the preference for energy substrates changes in the failing heart, there is less understanding about how this switch is governed.

Ca^2+^/calmodulin dependent protein kinase II (CAMK2) plays a crucial role in pathological cardiac remodeling induced by different pathological stimuli and its expression and activity are greatly increased in heart failure [9-15]. We have previously demonstrated that activation of CAMK2 through catecholaminergic stimulation of β-adrenergic receptors leads to phosphorylation of histone deacetylase 4 (HDAC4), thereby inducing its cytosolic accumulation and dissociation from the transcriptional activator myocyte enhancer factor 2 (MEF2) [16]. The dissociation of the MEF2-HDAC4 complex allows MEF2-dependent activation of a gene expression program that drives, among others, NR4A1-dependent activation of the hexosamine biosynthesis pathway (HBP), with downstream effects on protein O-linked glycosylation (O-GlcNAcylation) and calcium mishandling, causing HFrEF [17].

Adverse effects of CAMK2 also include altered mitochondrial calcium handling and oxidative phosphorylation [18], although CAMK2 does not control mitochondrial calcium handling under physiological conditions [19]. Other studies implicated CAMK2 as a regulator of cardiac metabolic remodeling [20-25]. One study demonstrated that CAMK2D deletion attenuates the maladaptive cardiac effects including mitochondrial dysfunction after pressure overload. It was suggested that CAMK2D-induced repression of peroxisome proliferator–activated receptor α (*Ppar-α*) and uncoupling protein 3 (*Ucp3*) is one of the drivers of mitochondrial dysfunction [25]. These findings suggest that CAMK2D is associated with cardiac metabolic remodeling during heart failure.

However, it remains unknown how CAMK2 regulates cardiac metabolic substrate switching leading to heart failure. In the current study, we establish CAMK2-NR4A1 signaling as an essential regulatory axis of pathological cardiac metabolic reprogramming. CAMK2, through upregulation of NR4A1, promotes substrate switching preceding HFrEF. We show that NR4A1 depletes the intracellular lipids reservoir in the myocardium by direct repression of the *Slc27a1* promoter, thereby preventing fatty acid uptake. Taken together, the data of this study establish a model by which metabolic substrate switching is initiated by the CAMK2-NR4A1 axis to directly induce lipid depletion, which is a trigger for the production of glucose by products that drive cardiac dysfunction. Thus, targeting cardiac lipid metabolism should become a major task to develop novel therapeutics for HFrEF.

## Results

### CAMK2-dependent induction of cardiac glucose uptake precedes HFrEF

The role of CAMK2 in HFrEF has been extensively investigated [9, 11, 12]. We previously demonstrated that the two cardiac CAMK2 isoforms (CAMK2D and CAMK2G) lead to HFrEF following pathological pressure overload [26]. Here, we utilized the newly established method of induction of pathological pressure overload named O-ring aortic banding (ORAB), which is compared to transverse aortic constriction more standardized and presents with less inter-surgeon variability [27]. Consistent with our previous work, echocardiographic, histological and gravimetric analysis revealed that cDKO mice are protected from HFrEF and fibrosis (**Supl. Fig. 1A-F**). Based on a previous report showing a continuous increase in cardiac glucose uptake from the first day up to 4 weeks in mice subjected to pressure overload [28], we now investigated whether this depends on cardiac CAMK2. Because we asked whether these metabolic changes may be causative for HFrEF, we conducted FDG-PET measurements before systolic dysfunction was established. Strikingly, we detected upon ORAB a robust increase in myocardial glucose uptake in Cre^+^ (served as controls) but not cDKO, preceding HFrEF (**Fig. 1A**). The quantification of dynamic glucose uptake revealed a 2-fold higher myocardial glucose uptake in Cre^+^ versus cDKO mice (**Fig. 1-B-C**). We then stimulated adult mouse ventricular myocytes (AMVMs) isolated from cDKO and Cre^+^ mice for two hours with the non-selective β-adrenergic agonist isoproterenol (ISO) and measured glucose uptake using 2-NBDG. Again, glucose uptake was robustly increased in AMVMs from Cre^+^ but not cDKO (**Fig. 1D**). These data indicate an essential role of CAMK2 in cardiac glucose uptake after pathological pressure overload in vivo and catecholaminergic stimulation in vitro.

**Figure 1.**
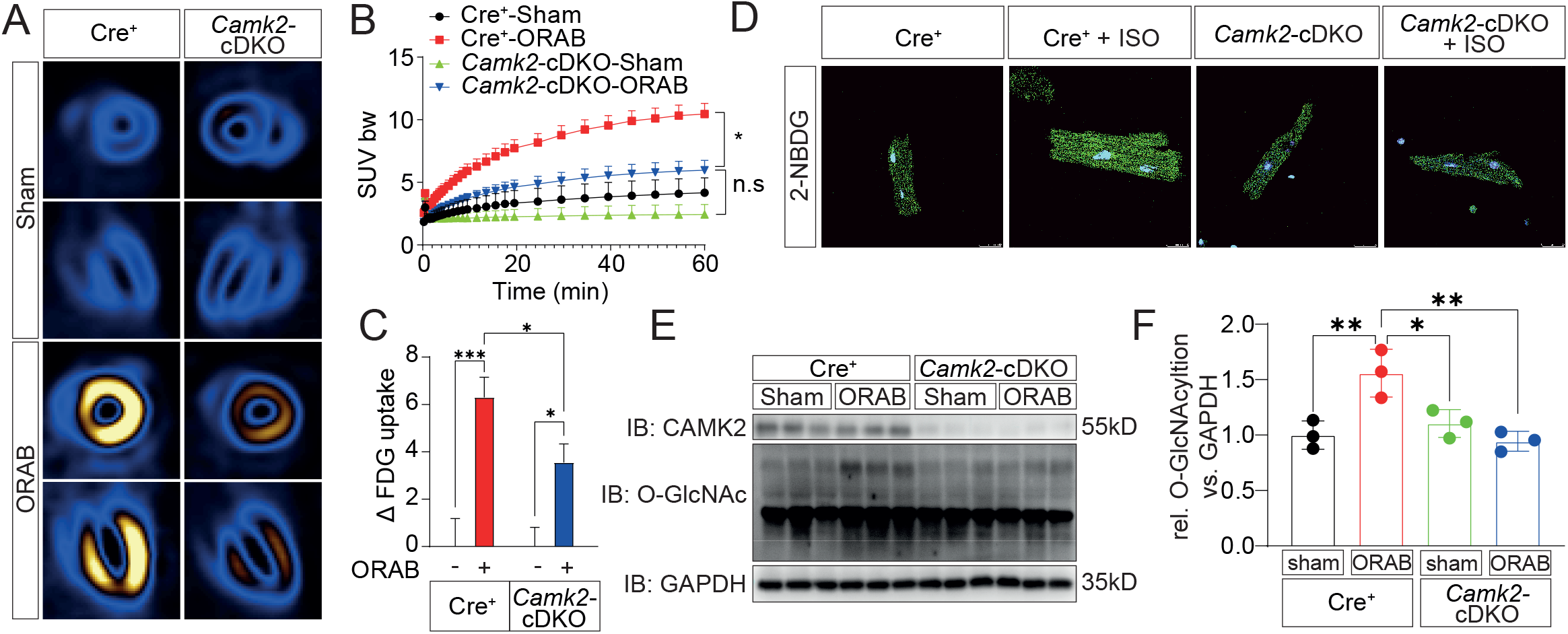
CAMK2 is involved in cardiac glucose uptake following ORAB-induced pressure overload. **A)** Myocardial glucose uptake using dynamic positron emission tomography (FDG-PET) reveals a robust increase in cardiac glucose uptake in Cre^+^ mice after ORAB-induced pressure overload compared to cDKO littermates. **B-C)** Quantification of FDG uptake during 60 min and analysis of delta values of FDG uptake after ORAB in Cre^+^ and cDKO littermates. (n=6). **D)** Representative fluorescent images of 2-NBDG uptake in isolated AMVMs obtained from Cre^+^ mice and cDKO littermates and stimulation with isoproterenol (ISO) for 2 hours. Scale bar is 20 μm. n = 10. **E)** Cardiac lysates from Cre^+^ mice as well as cDKO littermates two weeks after induction of pressure overload (ORAB) or sham surgery were immunoblotted (IB) with antibodies directed against CAMK2 and total protein O-GlcNAcylation. **F)** Quantification of immunoblot analysis showed upregulation of protein O-GlcNAcylation in the cardiac tissues of the Cre^+^ mice after ORAB compared to cDKO littermates (n = 3 mice/group). GAPDH was used as a loading control.

Although glycolysis and glucose oxidation in mitochondria is the canonical fate of glucose uptake, recent studies highlighted the importance of glucose side pathways including the HBP, as an alternative fate of glucose utilization. The HBP metabolizes glucose to an intermediate as a substrate for protein O-GlcNAcylation [17, 29, 30]. Since cardiac glucose uptake is increased in Cre^+^ mice following pathological pressure overload as compared to cDKO, we hypothesized that protein O-GlcNAcylation might be consequently activated on a CAMK2-dependent manner. Therefore, we determined the overall protein O-GlcNAcylation levels. Strikingly, overall protein O-GlcNAcylation was markedly increased in Cre^+^ but not cDKO mice (**Fig. 1E-F**), revealing that CAMK2 promotes a relative increase in glucose diversion into the HBP and protein O-GlcNAcylation. In the remaining manuscript, protein O-GlcNAcylation will serve as a surrogate parameter for pathological glucose flux.

### Downregulation of lipid metabolism as a very early event may regulate glucose metabolism

As mentioned previously, the failing myocardium is generally characterized by altered substrate utilization including a shift from FA to glucose utilization [2, 3, 31]. To gain further insights into the mechanistic order of events occurring during the metabolic substrate switching and whether these events precede the onset of HFrEF, we induced pressure overload in mice using ORAB for 1, 7 and 21 days. RNA sequencing analysis of left ventricular myocardium revealed a continuous decrease in expression of FA metabolic genes from the first day up to 21 days in mice subjected to ORAB surgery. Although, there were no appreciable changes in the expression of the genes involved in glucose metabolism 1 day after ORAB, we detected a significant upregulation of glycolytic and HBP genes from 7 days after ORAB, which indicates that downregulation of FA metabolism is an earlier event which may trigger the activation of glucose metabolism secondarily and all these precede HFrEF (**Fig. 2A-C**).

**Figure 2.**
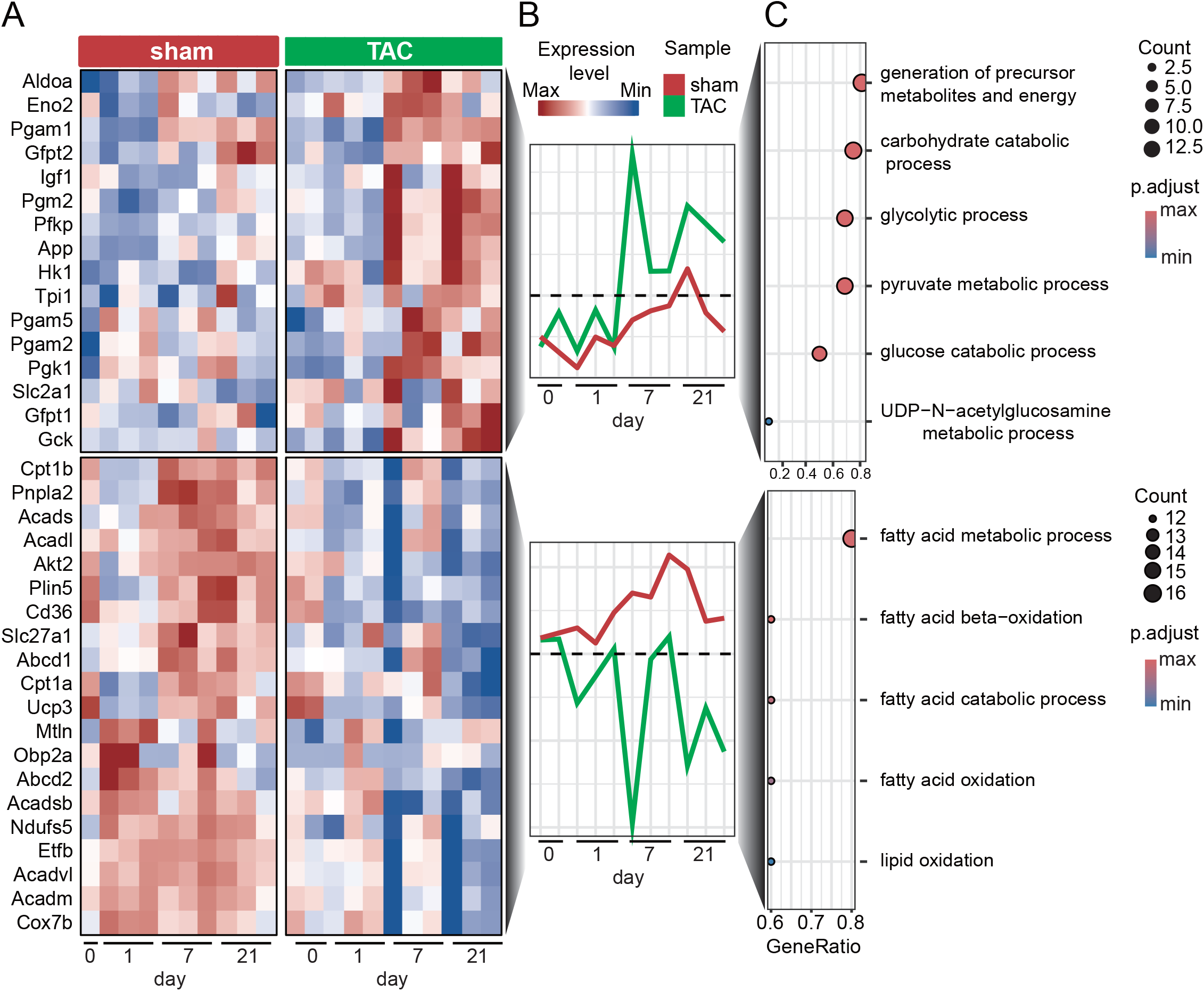
Identification of metabolic substrate switching as an early event following ORAB-induced pressure overload. **A)** RNA sequencing of ventricular myocardial tissues obtained from C57BL/6N wild type mice (WT) at 1, 7 and 21 day after ORAB-induced pressure overload. Heatmap depicting the expression levels of genes associated with the glycolytic process pathway (upper panel) and FA oxidation pathway (lower panel). **B)** Quantification of expression levels of the genes shown in heatmap, representing the median values across each time point for all involved genes. **C)** Gene Ontology (GO) term enrichment analysis highlights functional annotations of the pathway-related genes.

### Cardiac CAMK2 is required for lipid depletion in the myocardium

We applied deep RNA sequencing to uncover genes that were differentially regulated in the hearts of Cre^+^ mice subjected to pressure overload compared with cDKO mice. RNA sequencing identified 1273 differentially CAMK2-dependent expressed genes (**Fig. 3A**). Pathway enrichment and gene ontology analysis further revealed that a number of pathways involved in mitochondrial oxidative phosphorylation, adipogenesis and FA metabolism were among the most downregulated pathways in Cre^+^ mice after pressure overload (**Fig. 3B**). To further confirm that CAMK2 suppresses FA metabolism, cardiac lipid content in the heart was assessed by lipid droplet staining. The results showed a depletion of intramyocardial lipid reservoirs in Cre^+^ mice after pressure overload compared to Sham, which was prevented in cDKO littermates (**Fig. 3C)**. In addition, we performed steady-state lipidome analysis on cardiac samples. The results also revealed depletion of storage lipids triacylglycerols (TAGs) in cardiac tissues obtained from Cre^+^ as compared to cDKO mice after pressure overload (**Supl. Fig. 2A**). Furthermore, AMVMs isolated from cDKO and Cre^+^ mice were stimulated with ISO for two hours and subsequently intracellular lipid droplet content was detected using BODIPY staining. Indeed, intracellular lipid droplet contents were markedly higher in cDKO as compared to Cre^+^ AMVMs after ISO treatment (**Fig. 3D**). Transcriptome analysis revealed that nuclear receptor subfamily 4 group A member 1 (*Nr4a1*), a metabolic regulator and known driver of the HBP [17], was robustly upregulated in Cre^+^ but not cDKO mice. In stark contrast, genes involved in FA uptake and oxidation were significantly downregulated. For instance, *Slc27a1* (encoding for the FA transporter 1, FATP1), *Ucp3*, and components of the mitochondrial electron transport chain (such as *Etfb*) were markedly downregulated (**Fig. 3E**). As a sign of cardiac metabolic substrate switching, we observed activation of glycolytic genes and suppression of FA oxidation genes in Cre^+^ mice after pressure overload compared to cDKO (**Supl. Fig. 2B**). Together, these data indicate an essential role of cardiac CAMK2 in the regulation of FA metabolism after pathological pressure overload *in vivo* and catecholaminergic stimulation *in vitro*.

**Figure 3.**
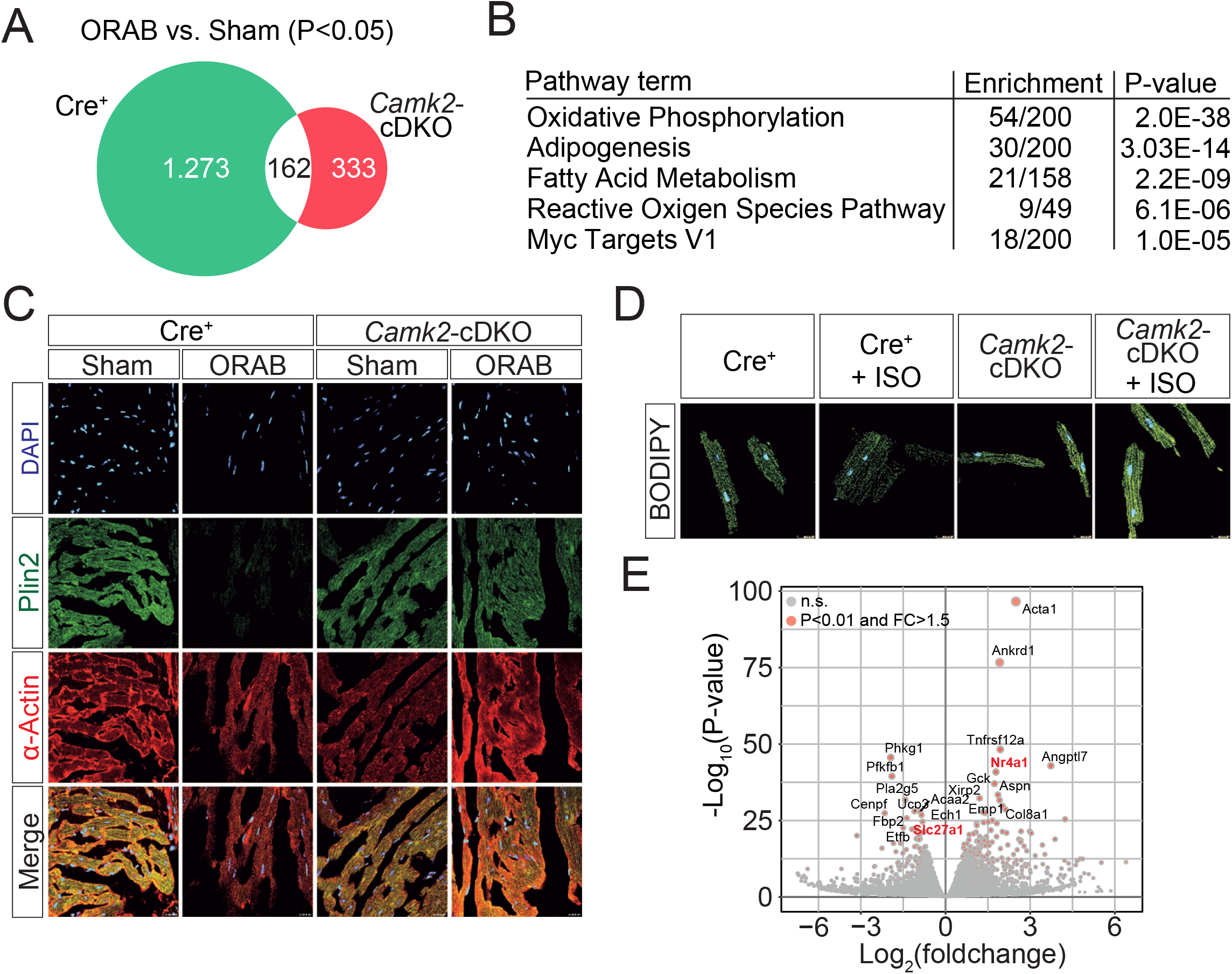
Cardiac CAMK2 regulates lipid metabolism. **A)** RNA sequencing of cardiac tissues obtained from Cre^+^ mice and cDKO littermates two weeks after ORAB-induced pressure overload. Venn diagram shows a total 1273 CAMK2-dependent differentially regulated genes. **B)** Pathway enrichment reveals top 5 *CAMK2*-dependent biological processes which are significantly downregulated after pathological pressure overload. **C)** Immunohistochemistry for the staining of lipid droplets shows drastic depletion of lipid droplet reservoir in Cre^+^ mice after ORAB-induced pressure overload as compared to cDKO. LV sections stained with Plin2 to visualize lipid droplets (green), a-actinin to visualize myocardium (red) and 4⍰,6-diamidino-2-phenylindole for nuclei staining (DAPI, blue). Scale bar is 20 μm. **D)** Representative fluorescent images of BODIPY staining for lipid droplets in isolated AMVMs obtained from Cre^+^ mice and cDKO littermates and stimulation with ISO for 2 hours. Scale bar is 20 μm. n = 10. **E)** Volcano plot shows the most robust *CAMK2*-dependent target genes in ORAB compared to Sham groups, e.g. *Nr4a1* and FA transporter *Slc27a1* highlighted in red. Regulated mRNAs were significantly considered at a fold change of FC >1.5 and P<0.01.

### CAMK2 is sufficient to induce metabolic substrate switching

Next, we tested whether adeno-associated virus (AAV) mediated overexpression of *Camk2* is sufficient to induce metabolic substrate switching. We overexpressed the predominant cardiac CAMK2D-C isoform in a constitutive active form (Myc-tagged *Camk2D-C*-T287D) specifically in cardiomyocytes by injection of 1×10^12^ virus particles under the control of a cardiac-specific troponin-T promoter in cardiotrophic AAV9 (AAV9-*Camk2D-C*). The firefly luciferase gene was cloned into the same AAV9 vector to generate control particles (AAV9-Luc), and either AAV9-*Camk2D-C* or AAV9-Luc was injected into C57BL/6N mice (**Fig. 4A**). Four weeks after viral transduction, we observed, as compared with control AAV9-Luc mice, a severe HFrEF phenotype, as well as increased fibrosis as assessed by staining of the extracellular matrix by trichrome (**Supl. Fig. 3A-D**). We confirmed overexpression of *Camk2D-C* as well as upregulation of *Col1a1, Nppa* and *Nppb* (**Supl. Fig. 3E-H**). These findings are consistent with prior work by Zhang et al. (2003), who demonstrated that activation of the *Camk2D-C* isoform promotes cardiac hypertrophy, dilated cardiomyopathy, and heart failure [32].

**Figure 4.**
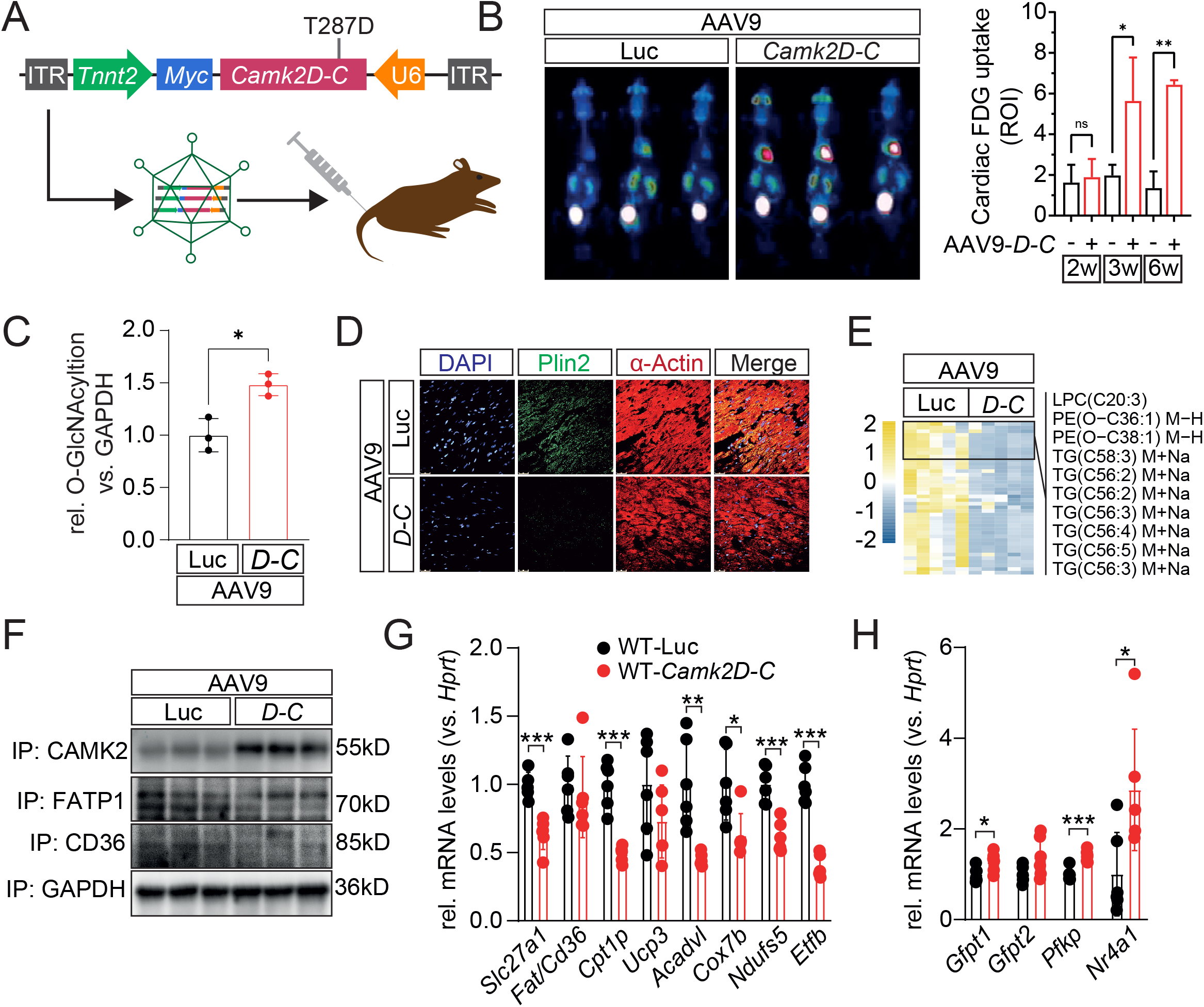
Cardiac overexpression of CAMK2D-C induces metabolic remodeling. **A)** Scheme of the production and gene delivery approach of cardio trophic AAV9-Luc (control vector) and AAV9-*Camk2D-C* constitutive active *T287D* form and injection to the mice. **B)** Myocardial glucose uptake using dynamic positron emission tomography (FDG-PET scanning) reveals a robust increase in cardiac glucose uptake in mice overexpressing AAV9-Camk2*D-C* compared to AAV9-Luc control littermates. Quantification of FDG uptake during 60 min is shown. **C)** Quantification of immunoblot analysis of total protein O-GlcNAcylation shown from cardiac tissues obtained from AAV9-*Camk2D-C* as well as AAV9-Luc control littermates (n = 3 mice/group). **D)** Immunohistochemistry for the staining of lipid droplets shows drastic depletion of lipid reservoir in the mice overexpressing AAV9-*Camk2D-C* compared to AAV9-Luc control littermates. Left ventricular (LV) sections stained with Plin2 to visualize lipid droplets (green), a-actinin to visualize myocardium (red) and for nuclei staining (DAPI, blue). Scale bar is 20 μm. **E)** Heatmap of steady-state metabolomic analysis reveals depletion of TAGs in the mice overexpressing AAV9-*Camk2D-C* compared to AAV9-Luc control littermates (n=5). **F)** Cardiac lysates from mice overexpressing AAV9-*Camk2D-C* as well as AAV9-Luc control littermates were immunoblotted (IB) with antibodies directed against *CAMK2*, FATP1, and FAT/CD36. Representative immunoblot analysis showed downregulation of fatty acid transporters FATP1 and FAT/CD36 in the cardiac tissues of the mice overexpressing CAMK2D-C (*n* = 3 mice/group). GAPDH was used as a loading control. **G-H)** Gene expression analysis of downregulation of FA metabolic pathway and upregulation of HBP-related genes. All values are presented as mean ± SEM. \**P*<0.05. n.s. indicates not significant.

To further investigate whether overexpression of CAMK2D-C is sufficient to induce myocardial glucose uptake, we measured myocardial glucose uptake using FDG-PET scanning of mice. Strikingly, we observed a significant increase of myocardial glucose uptake in mice overexpressing AAV9-*Camk2D-C* as compared to AAV9-Luc control littermates in a time course of 2 – 6 weeks after viral transduction (**Fig. 4B**). Accumulation of protein O-GlcNAcylation in cardiac samples obtained from AAV9-*Camk2D-C* further confirmed increased glucose flux (**Fig. 4C**). Moreover, we detected elevation of UDP-N-acetylglucosamine (UDP-GlcNAc) in water soluble metabolites in cardiac extracts of AAV9-*Camk2D-C* mice compared to AAV9-Luc (**Supl. Fig. 3I**).

Lipid droplet staining by Plin2 immunohistochemistry showed a depletion of intramyocardial lipid reservoirs in the AAV9-*Camk2D-C* mice as compared to AAV9-Luc (**Fig. 4D**). In addition, we performed steady-state lipidomics analysis on cardiac samples from AAV9-*Camk2D-C* or AAV9-Luc. The results revealed a substantial depletion of storage lipids TAGs in cardiac tissues obtained from AAV9-*Camk2D-C* as compared to AAV9-Luc (**Fig. 4E**). TAGs have been identified as significant sources of fuel for mitochondrial oxidation in the heart [33].

Next, we tested whether CAMK2D-C induced cardiac lipid depletion is due to impaired FA import or excessive utilization. Notably, overexpression of *Camk2D-C* decreased levels of the two cardiac FA transporters FATP1 and CD36 (**Fig. 4F**), indicating that impaired FA uptake might be the cause of the cardiac lipid depletion. To further validate the CAMK2-targeted genes obtained from RNA-Seq analysis, we analyzed the expression levels of the genes involved in FA uptake and oxidation along with the HBP-related genes. Interestingly, we found an overall suppression of FA uptake and oxidation genes in AAV9-*Camk2D-C* mice, which was associated with significant upregulation of *Nr4a1* and other HBP-related genes (*Gfpt1, Gfpt2*) (**Fig. 4G-H**). Taken together, these data indicate that AAV9-mediated overexpression of *Camk2D-C* regulates cardiac metabolic remodeling on one hand by induction of cardiac glucose uptake and activation of the HBP pathway; and on the other hand by repression of FA uptake and oxidation.

### NR4A1 acts downstream of CAMK2

One of the gene transcripts that was found to depend on cardiac CAMK2 and to be induced by CAMK2 is *Nr4a1* (**Fig. 3E and 4H**). Evidence has emerged that NR4A1 plays an important role in cardiac metabolism and in particular in controlling glycolytic flux and oxidative stress. Our lab previously proposed that Nr4A1 mediates activation of the HBP [17], and other studies were able to confirm these findings [34-37]. However, until today the exact underlying mechanism of this regulation has not been elucidated and the role of Nr4A1 has not been studied *in vivo*. Here, we generated inducible cardiomyocyte-specific *Nr4a1* knock-out (*Nr4a1-*cKO) mice. Cre-positive mice in the same genetic background served as controls. The Cre^+^ and *Nr4a1*-cKO mice were treated with tamoxifen for two weeks and subsequently subjected to ORAB surgery to induce pathological pressure overload (**Fig. 5A-B**). The efficiency of inducible knock-out was confirmed by analyzing *Nr4a1* expression using RT-qPCR (**Fig. 5C**). Four weeks after ORAB-induced pressure overload, the hearts of *Nr4a1*-cKO mice were protected from HFrEF, without impacting heart weight to body weight ratios (**Fig. 5D-E**). Gene expression analysis of *Col1a1* and trichrome staining of transverse section of heart tissues revealed less fibrosis in *Nr4a1*-cKO mice following pressure overload as compared to Cre^+^ littermates (**Supl. Fig. 4A-C)**. Thus, deficiency of *Nr4a1* in cardiomyocytes protects the heart from functional and structural remodeling after ORAB-induced pressure overload.

**Figure 5.**
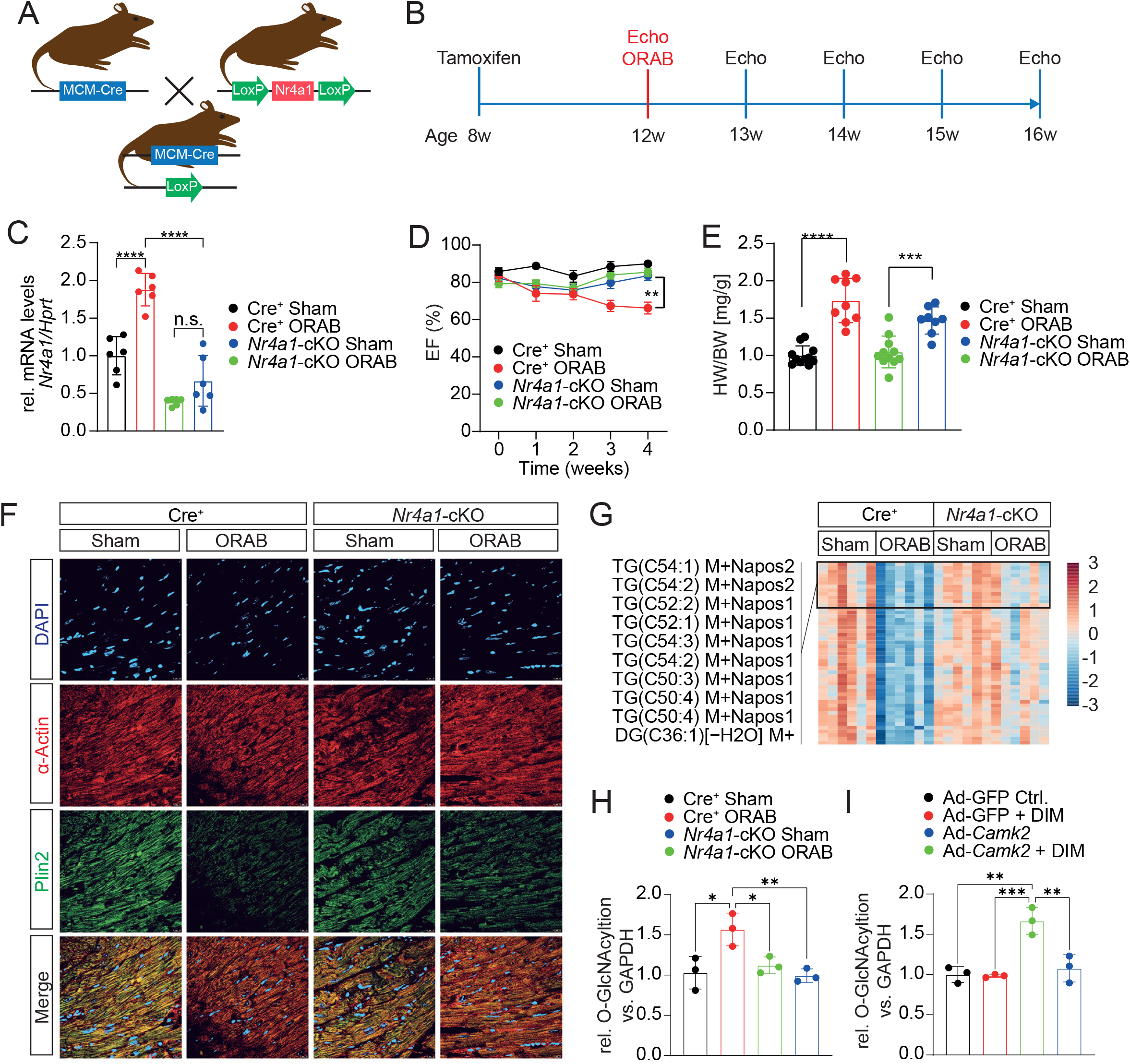
*Nr4a1*-cKO attenuates cardiac dysfunction and lipid depletion following ORAB-induced pressure overload. **A-B)** Scheme of generation of inducible cardiomyocyte-specific *Nr4a1* knockout (KO) mice. *Nr4a1*-cKO and Cre^+^ littermate mice were randomized to either ORAB or sham surgery and euthanized after 4 weeks. **C)** Fold changes in mRNA levels of *Nr4a1*. All values are presented as mean ± SEM. \**P*<0.05. n.s. indicates not significant. **D)** Values of left ventricular ejection fraction and **E)** quantification of heart weight/body weight ratios are shown (n≥8 per group). **F)** Immunohistochemistry for the staining of lipid droplets shows drastic depletion of lipid droplet reservoir in Cre^+^ mice after ORAB-induced pressure overload as compared to *Nr4a1*-cKO. LV sections stained with Plin2 to visualize lipid droplets (green), a-actinin to visualize myocardium (red) and for nuclei staining (DAPI, blue). Scale bar is 20 μm. **G)** Heatmap of steady-state lipidomic analysis revealed depletion of TAGs in Cre^+^ mice after ORAB-induced pressure overload as compared to *Nr4a1*-cKO littermates. Significantly regulated lipids (one-way ANOVA with Tukey post-test) are shown. (n=6). **H)** Quantification of immunoblot analysis of total protein O-GlcNAcylation shown from cardiac tissues obtained from Cre^+^ mice as well as *Nr4a1*-cKO littermates four weeks after induction of pressure overload (n = 3 mice/group). **I)** Human iPS-CMs were adenovirally infected with either constitutively active *Camk2D-C* or GFP vector as a control for 24 hours and then incubated with or without the NR4A1 inhibitor DIM-C-p-PhOH (DIM) for next 24 hours. Quantification of immunoblot analysis of total protein O-GlcNAcylation is shown (*n* = 3 cultures/group). All values are presented as mean ± SEM. \**P*<0.05. n.s. indicates not significant.

Several studies have shown a role of NR4A1 in glucose and FA metabolism [38-42]. Thus, we hypothesized that *Nr4a1* as a CAMK2-downstream gene might mediate metabolic substrate switching. To determine whether *Nr4a1* deletion alters lipid content in the heart after pressure overload, lipid droplet staining in cardiac tissue was performed. The results showed a depletion of the intramyocardial lipid reservoir in Cre^+^ mice after pressure overload, which was blunted by *Nr4a1*-cKO (**Fig. 5F**). Steady-state lipidome analysis. revealed depletion of TAG contents in Cre^+^ as compared to *Nr4a1*-cKO mice (**Fig. 5G**).

Consistent with previous *in vitro* findings [17], deficiency *of Nr4a1 in vivo* blunted the increased levels of O-GlcNAcylation after ORAB, which phenocopied our observation in cDKO mice (**Fig. 5H**). To prove that CAMK2 activates the HBP and protein O-GlcNAcylation levels through *Nr4a1*, we overexpressed constitutive active CAMK2D (T287D) in human iPS-CMs. Increased protein O-GlcNAcylation levels were blunted by the NR4A1 inhibitor DIM (**Fig. 5I**), formally confirming that NR4A1 acts downstream of CAMK2 to induce metabolic substrate switching.

### NR4A1 suppresses the FA transporters FATP1 and CD36

At the mRNA levels, adenoviral overexpression of *Nr4a1* in human iPSC-CMs led to a significant down-regulation of the FA transporters FATP1 (*Slc27a1*), *Cd36* and mitochondrial genes involved in FA oxidation (such as *Acadvl*), whereas genes encoding for proteins for glucose utilization were upregulated (e.g., *Slc2a1, Gfpt2*) (**Fig. 6A**). We also analyzed the same genes in myocardial samples from explanted hearts of patients with heart failure compared with nonfailing donors (from healthy donor hearts that could not be transplanted), and we found a similar regulation of *SLC27A1, CD36* and other enzymes involved in β-oxidation, whereas *NR4A1* was strongly upregulated (**Fig. 6B**). Thus, our data derived from experimental NR4A1 overexpression strongly correlate with the metabolic state in end-stage human heart failure.

**Figure 6.**
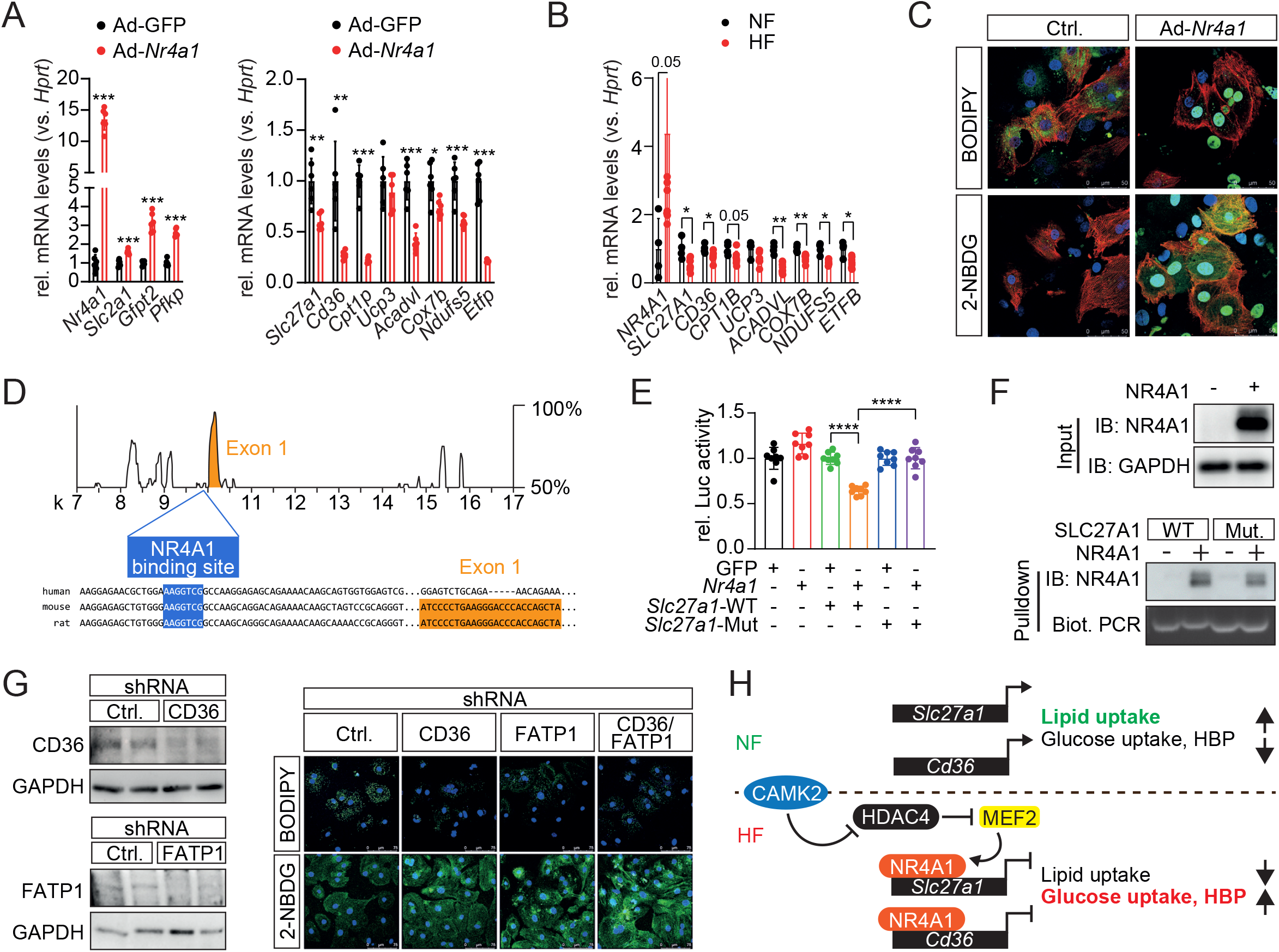
NR4A1 directly binds and suppresses FA transporter FATP1 gene expression. **A)** Gene expression analysis in human iPS-CM, as assessed by real-time PCR, 48 hours after transduction with adenovirus encoding either GFP-tagged *Nr4a1* (Ad-*Nr4a1*) or GFP empty vector as a control (Ad-GFP) (*n* = 6 cultures/group). **B)** Gene expression analysis measured by qRT-PCR confirmed upregulation of *Nr4a1* and downregulation of FA uptake and FA oxidative machinery in patients with failing heart (HF, n=10) as compared to non-failing heart (NF, n=5). All values are presented as mean ± SEM. \**P*<0.05. n.s. indicates not significant. **C)** Detection of lipid content and glucose uptake using BODIPY and 2-NBDG staining 48 hours after NR4A1 overexpression in cardiomyocytes. **D)** In silico analysis illustrated putative NBRE-like sequence as potential NR4A1 binding site on the promoter of FA transporter FATP1 (*Slc27a1*), which is a conserved region in human, mouse and rat species. **E)** Effect of NR4A1 on transcriptional activities of the wild type (WT) and a mutated sequence of *Slc27a1* promoters, detected by luciferase reporter assay in C_2_C_12_ cells (*n* = 8 cultures/group). All values are presented as mean ± SEM. \**P*<0.05. n.s. indicates not significant. **F)** The occupation of NR4A1 on the NBRE-L sequence in the promoter of *Slc27a1* was assessed by a biotinylated DNA-Protein pull down assay. **G)** The protein expression levels of FATP1 and CD36 in human iPS-CM, 48 hours after lentiviral knockdown assay detected by western blotting. GAPDH was used as a loading control. Detection of lipid content and glucose uptake using BODIPY and 2-NBDG staining 48 hours after lentiviral knockdown assay in human iPS-CM. **H)** Schematic model of action shows how *CAMK2*-NR4A1 mediates metabolic substrate switching.

To determine whether NR4A1-induced down-regulation of FA transporters affects cellular lipid content and glucose uptake, we utilized BODIPY and 2-NBDG staining, respectively. While lipid content was decreased in cardiomyocytes overexpressing NR4A1, glucose uptake was increased, providing direct evidence for a role of NR4A1 in metabolic substrate switching (**Fig. 6C**). To gain more insights into the mechanism underlying the NR4A1-dependent repression of FA transporters FATP1 and FAT/CD36, we tested whether NR4A1, which is mainly located in the nucleus, may function as a transcriptional repressor to control FA uptake. In silico sequence analysis using the VISTA software revealed in the *Slc27a1* promoter a NBRE-like (NBRE-L) sequence that has been described as a response element for NR4A1 [34, 40]. This sequence is conserved in human, mouse and rat (**Fig. 6D**). A similar response element for *Nr4a1* was previously described in the CD36 promoter [40]. Luciferase reporter assay revealed that a mutation of this NBRE-L element was sufficient to abrogate the suppressive function of NR4A1 on *Slc27a1* promoter activity (**Fig. 6E**), confirming functional relevance. We then amplified a biotinylated fragment of the *Slc27a1* promoter region containing the NR4A1-binding site or a mutated NR4A1-binding site. In a pulldown assay, we identified a direct interaction between NR4A1 and the NBRE-L sequence in the *Slc27a1* promoter (**Fig. 6F**). These data indicate that NR4A1 binds to the *Slc27a1* promoter, which leads to repression of its transcription. To investigate the functional role of FATP1 and CD36 in lipid uptake and determine whether impaired FA uptake causes cardiac lipid depletion, FATP1 and CD36 were separately or simultaneously knocked down using lentiviruses containing small hairpin (sh) RNAs. 48 hours after lentiviral infection of human iPSC-CMs, we measured the lipid content and glucose uptake. Collectively, our data showed that suppression of FA transporters FATP1 and CD36 elicits a reduction in lipid uptake. Strikingly, knockdown of both transporters triggered elevated glucose uptake in a synergistic manner, indicating that alterations of FA transporter can per se impact glucose metabolism (**Fig. 6G**). Taken together, our data show that NR4A1 directly represses FA uptake by transcriptional repression of both FA transporters on cardiomyocytes which subsequently leads to an induction of glucose uptake.

## Discussion

The pathogenesis of HFrEF is marked by metabolic reprogramming to allow shifting from FA to glucose utilization and activation of glucose-derived side pathways such as the HBP [2, 29, 43]. The present study provides clear evidence that CAMK2 induces metabolic substrate switch from FA to glucose utilization by inducing the expression of *Nr4a1*. NR4A1 binds to the promoters of the FA transporters *Cd36* and *Slc27a1* to repress their expression. Reduction in FA uptake is sufficient to induce glucose utilization and HBP (**Fig. 6H**).

Metabolic alterations play a pivotal role in the pathogenesis of HFrEF, and understanding the temporal sequence of these events provides critical insights into the underlying mechanisms of disease progression. Here, we could precisely observe down-regulation of FA-related genes at an early stage during the development of pressure overload induced HF, which was followed by the activation of glucose metabolic genes as a compensatory mechanism. This shift in gene expression profiles from FA towards glucose metabolism underscores the dynamic nature of metabolic adaptation of the heart in response to pathological stimuli. Crucially, this shift in gene expression, indicative of metabolic substrate switching, precedes the onset of HFrEF. Unraveling these early molecular events provides a window of opportunity for preventive strategies and for reversal of disease by targeting the underlying driving force. This insight also prompts further investigations into the causative factors triggering metabolic remodeling, offering potential therapeutic targets to delay or prevent the progression to advanced stages of HF.

It is generally acknowledged that decreased expression and activity of proteins involved in cardiac FA uptake and oxidation lead to impaired mitochondrial function in advanced heart failure [5]. It has been reported that CAMK2 activation induces mitochondrial reprogramming through down-regulation of transcription factor *Ppar-α* and mitochondrial protein *Ucp3* following pressure overload [25]. NR4A1 has also been shown to play a role through down-regulation of *Pparγ2*, a key regulator of FA metabolism, in white adipose tissue [38]. However, additional studies must be performed to verify the connection between the CAMK2-NR4A1 regulatory axis with PPARs signaling mechanisms in the regulation of FA metabolism in the heart.

NR4A1 was found to be upregulated after pathological cardiac stress [36] and is directly regulated by the myocyte enhancer factor 2 (MEF2) transcription factor [44]. A few studies started to study the underlying mechanisms of NR4A1 in mediating cardiac pathologies [34]. It has been suggested that NR4A1 positively regulates cardiac hypertrophy and Ca^2+^ homeostasis [45]. Our previous study showed that NR4A1 drives the expression of *Gfpt2* and induces the HBP and aberrant cardiac protein O-GlcNAcylation [17]. Here, we used for the first time an inducible deletion of *Nr4a1* in cardiomyocytes and could confirm a maladaptive role of NR4A1 due to repression of FA uptake. However, NR4A1 plays an important role in immune cells where it seems to play an anti-inflammatory role [46-48]. The contrasting roles of NR4A1 in cardiomyocytes and immune cells highlight its adaptability to different cellular contexts. Understanding the distinct roles of NR4A1 in these cell types is crucial for developing targeted therapeutic interventions, especially in conditions where both cardiac metabolism and immune responses are involved, such as in certain cardiovascular diseases or inflammatory conditions affecting the heart.

NR4A1 acts as a transcription factor driving the expression of several glucose metabolic genes [41] and as a repressor of FA metabolic genes [40] by binding to NBRE-L sequences in the promoter regions of genes. The dual role of NR4A1 as a transcriptional activator or suppressor is context-dependent and can be attributed to a combination of factors, including interactions with cofactors, binding partners and post-translational modifications [49]. The complexity of these interactions allows NR4A1 to play diverse roles in different biological processes and pathways. For instance, NR4A1 interacted with SWI/SNF corepressor complex on CD36 and FABP4 promoters to suppress their transcription. This impedes FA uptake leads to inhibition of cell proliferation in a breast cancer cell line [40]. In addition, NR4A1 inhibits expression of CD36 in macrophages [50] and white adipose tissue [38], suggesting a common role for NR4A1 in regulating lipid uptake in different tissues. Whether a similar suppressor complex regulates the expression of FA transporter CD36 in cardiomyocytes needs to be investigated. Furthermore, it needs to be tested whether a similar co-repressor complex might exist also for FATP1. Whereas the role of NR4A1 in other cell types, in particular immune cells is well established, its specific role in cardiomyocytes has just started to be dissected. Of note, we clearly identified NR4A1 as a maladaptive factor in cardiomyocytes. Notably, in immune cells it plays adaptive roles. Thus, context-dependent mechanisms and cell-type specific targeting strategies need to be developed.

The uptake of the majority of FA in the heart depends on several FA transporters, including FA transport proteins 1 and 6 (FATP1 and 6), FA translocase FAT/CD36 and plasma membrane associated FA binding protein (FABP) [51]. Dysregulation of FA transporter CD36 activity is implicated in metabolic disorders such as obesity, insulin resistance, or cardiovascular diseases [52]. In general, considerably more is known about the physiological and pathological regulation of CD36 compared to FATPs in the heart [51, 53, 54]. A few reports showed that cardiac-specific overexpression of FATP1 in mice leads to increase lipid accumulation and induced lipotoxic cardiomyopathy, along with a decreased rate of glucose utilization [55], while insulin-stimulated triacylglycerol synthesis was blunted in mice with FATP1 deficiency [56]. Here, we confirmed the functional relevance of FA transporters FATP1 and CD36 in FA uptake in cardiomyocytes using a knock-down approach. The observation that disruption of lipid metabolism triggers a compensatory increase in glucose uptake unveils a fascinating aspect of metabolic crosstalk within cardiomyocytes. This adaptive response underscores the metabolic plasticity of the heart, revealing its ability to dynamically adjust fuel preferences in response to perturbations in nutrient availability. This intricate interplay between lipid and glucose utilization suggests a finely tuned regulatory network that ensures the maintenance of energy homeostasis in cardiomyocytes. Moreover, this may provide insights into potential therapeutic strategies to re-direct lipid fluxes in an organ-specific fashion such as in the heart.

In line with our experimental finding in mice, we could also detect a strong correlation between CAMK2 and NR4A1 upregulation and decreased mRNA expression for selected enzymes of the FA uptake and oxidation pathway in myocardial samples of patients with end-stage heart failure, further emphasizing the potential for CAMK2-NR4A1 axis as a therapeutic target to restore metabolic balance. Moreover, increased NR4A1 expression levels indicate the potential for adverse cardiac remodeling and could serve as a prognostic marker for certain cardiac pathologies.

In conclusion, our data shed new light on the contribution of the CAMK2-NR4A1 axis to metabolic remodeling induced by pressure overload in the heart. We identify a decline in FA transporter FATP1 expression in response to pressure overload and demonstrate a distinct mechanism by which NR4A1 transcriptionally suppressed FA transporter FATP1, leading to impeded FA uptake. These findings lead to potential strategies for treating HFrEF through metabolic intervention.

## Acknowledgments

We thank Jutta Krebs-Haupenthal, Michaela Oestringer and Ina Broll for their technical support.

## Figure Legends

**Supl. Figure 1.**
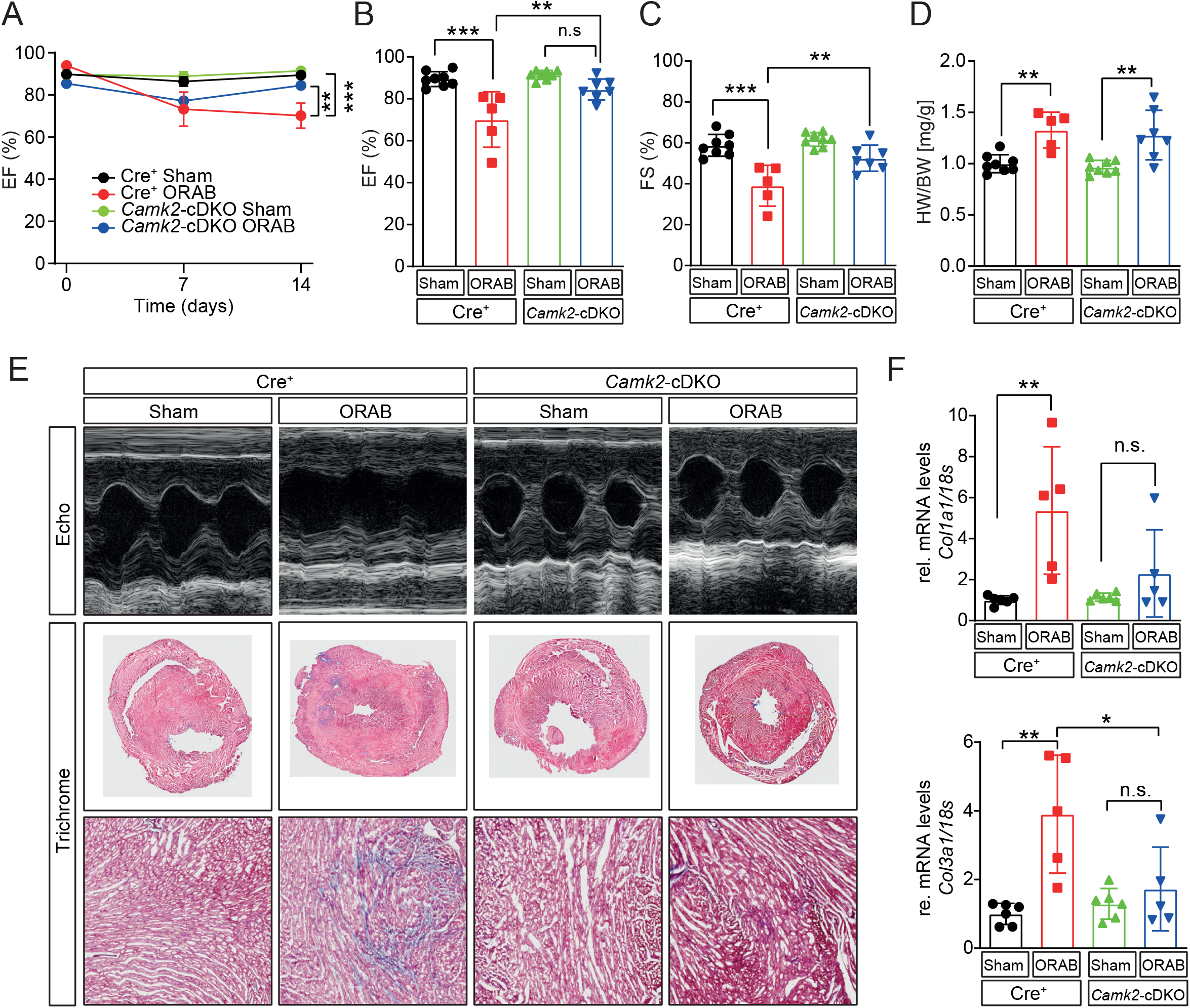
cDKO attenuates cardiac dysfunction following ORAB-induced pressure overload. cDKO and Cre^+^ (served as controls) littermate mice were randomized to either ORAB or sham surgery and euthanized after 2 weeks. **A-B)** Values of left ventricular ejection fraction, **C)** fractional shortening, and **D)** quantification of heart weight/body weight ratios are shown (n≥5 per group). **E)** Representative images of the echocardiographic M-modes, and Masson trichrome staining of transverse section of hearts are shown. **F)** Fold changes in collagen gene expression *Col1a1* and *Col3a1* as fibrotic markers. All values are presented as mean ± SEM. \**P*<0.05. n.s. indicates not significant.

**Supl. Figure 2.**
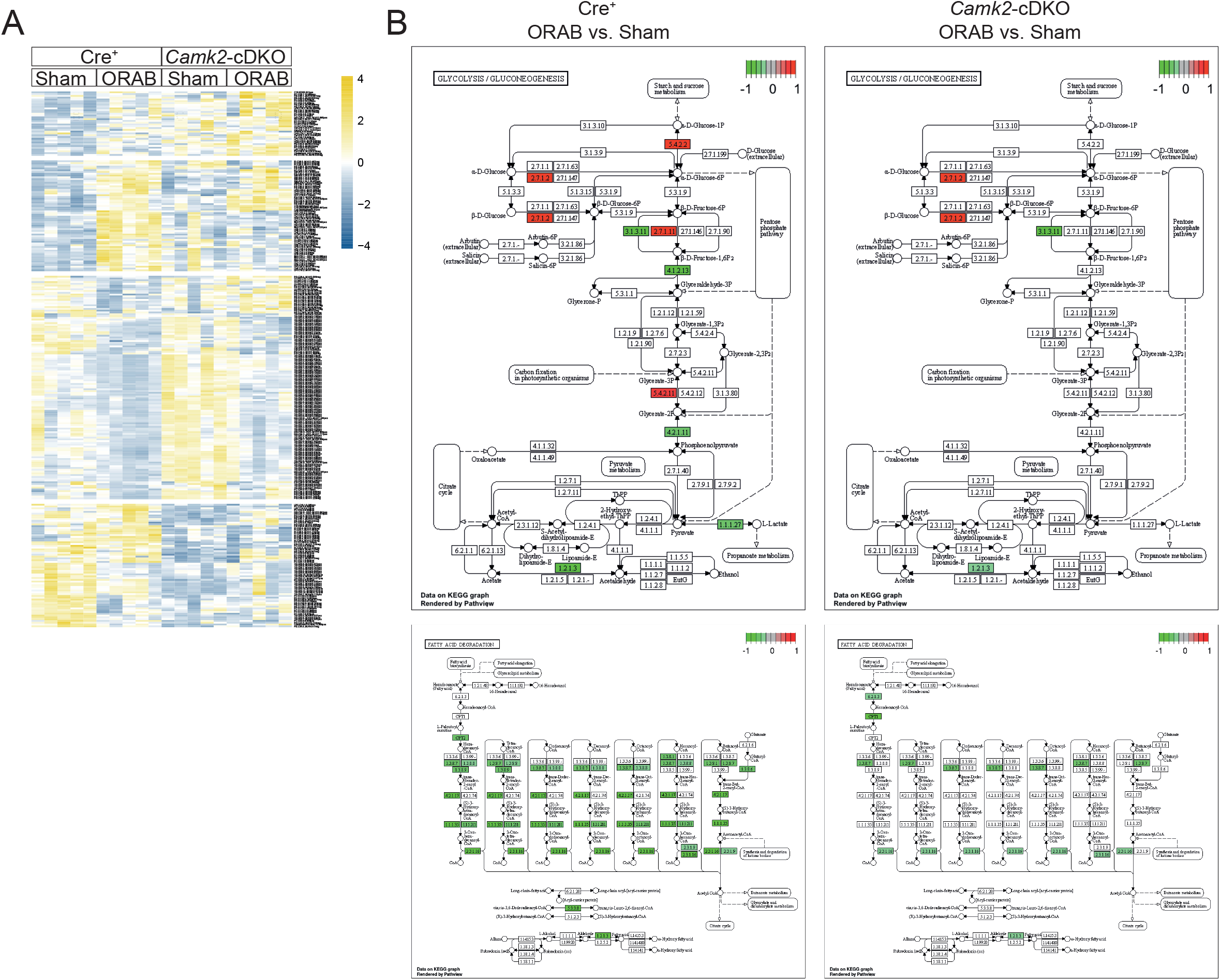
CAMK2 is involved in cardiac metabolic reprogramming. **A)** Heatmap of steady-state lipidomic analysis revealed depletion of triacylglycerols (TAGs) in Cre^+^ mice after ORAB-induced pressure overload as compared to cDKO littermates. (n=5). Blue indicates low expression; and yellow, high expression. **B)** Activation of glycolytic gene expression (top left panel) and suppression of FA oxidative gene expression in Cre^+^ mice after ORAB (bottom left panel) and cDKO (right panels) compare to corresponding sham control. KEGG pathway analysis was used. Green indicates low expression; and red, high expression.

**Supl. Figure 3.**
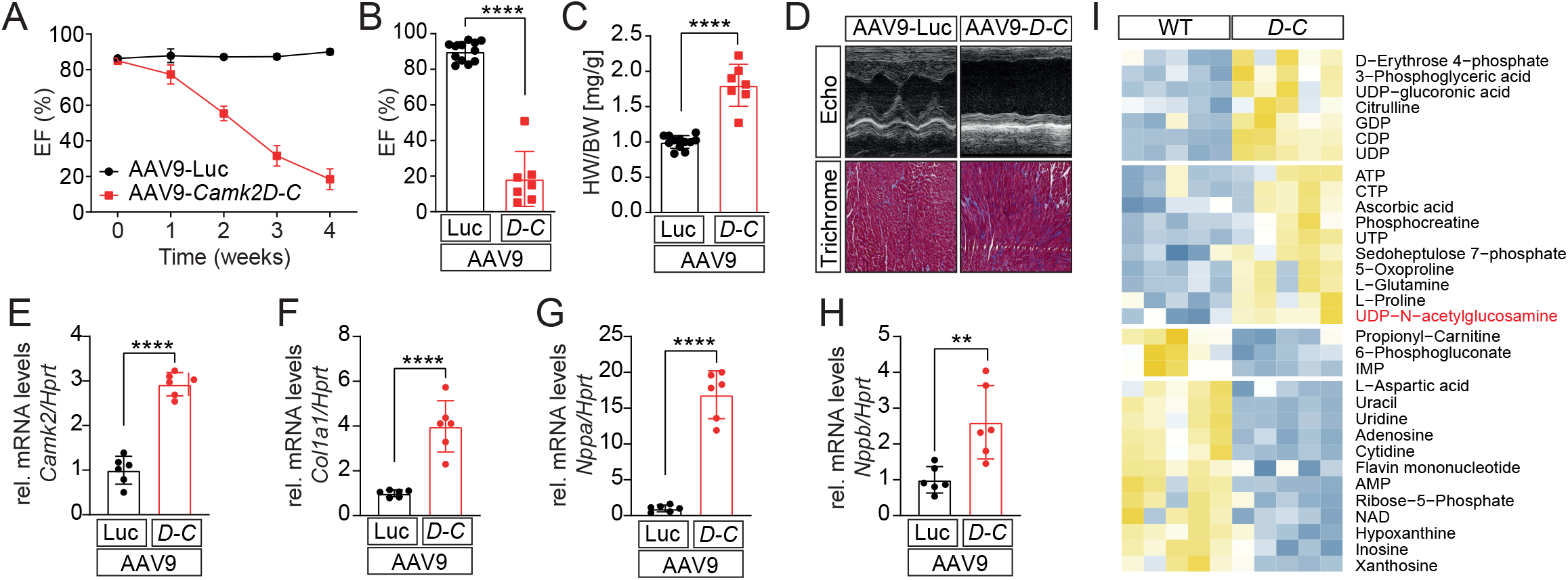
Cardiac overexpression of CAMK2D-C induces cardiac dysfunction. **A-B)** Time course of left ventricular ejection fraction and values of left ventricular ejection fraction and **C)** quantification of heart weight/body weight ratios are shown (n≥7 per group). **D)** Representative images of the echocardiographic M-modes and Masson trichrome staining of transverse section of hearts are shown. **E-H)** Fold changes in mRNA levels of *Camk2*, fibrotic marker collagen *Col1a1*, and hypertrophic markers *Nppa* and *Nppb*. All values are presented as mean ± SEM. \**P*<0.05. n.s. indicates not significant. **I)** Heatmap of steady-state water soluble metabolites profiling analysis in cardiac tissues obtained from AAV9-*Camk2D-C* and AAV9-Luc mice (n=5). Blue indicates low expression; and yellow, high expression.

**Supl. Figure 4.**
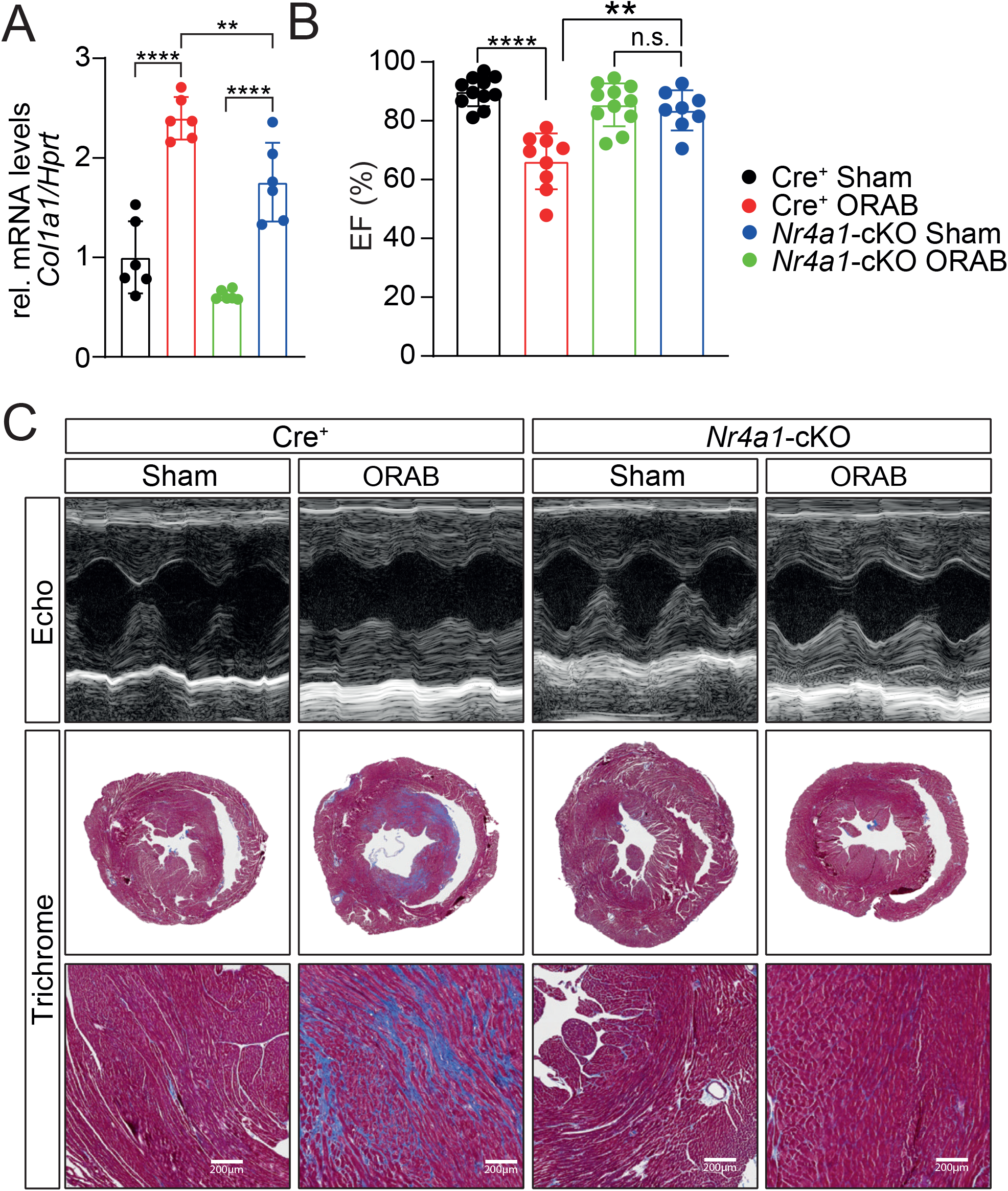
*Nr4a1*-cKO attenuates cardiac dysfunction and fibrosis following ORAB-induced pressure overload. **A)** Fold changes in mRNA levels of collagen gene *Col1a1* as fibrotic marker and **B)** values of left ventricular ejection fraction are shown. All values are presented as mean ± SEM. \**P*<0.05. n.s. indicates not significant. **C)** Representative images of the echocardiographic M-modes and Masson trichrome staining of transverse section of hearts are shown.

## Methods

### Human myocardial samples

Human myocardial samples were obtained from healthy donor hearts that could not be transplanted for technical reasons or from explanted hearts of patients with HF. Informed written consent was obtained from all donors and patients. The study conforms to the Declaration of Helsinki and was approved by the ethics committee of the University Medical Center Göttingen (Aktenzeichen 31/9/00).

### Mouse experiments

All experimental procedures were reviewed and approved by the Institutional Animal Care and Use Committee at the Regierungspräsidium Karlsruhe, Germany. We used conditional knockout mice containing cardiomyocyte-specific deletions of *Camk2d* and *Camk2g* (double knockout mice [cDKO]) as previously described [26]. To induce CAMK2D-C overexpression using AAV9, wild type mice with C57BL/6N background were used.

### Generation of *Nr4a1*-deficient mice

Inducible cardiomyocyte-specific *Nr4a1* knock-out (cKO) mice were generated by mating floxed *Nr4a1* mice with MerCreMer-transgenic mice under the control of the *NNT* promoter. MCM–Cre mice (Cre^+^) that were homozygous for the wild type (WT) *Nr4a1* allele were used as control. The mice were treated with tamoxifen to achieve *Nr4a1* deletion at 8 weeks of age. Tamoxifen (10 mg/ml, Sigma) was dissolved in a regular sunflower oil solution containing 6% ethanol. Tamoxifen (80 mg/Kg) was administered via gavage for 10 days (two days pause in between) to obtain cKO and Cre expressing control mice. Genotyping primer sequences were the following: Forward [TGACACCCTCACACGGACAA] and reverse [AGGACACCCATGCTCATGTG] to identify floxed *Nr4a1*, and forward [ATACCGGAGATCATGCAA GC] and reverse [AGGTGGACCTGATCATGGAG] to genotype the Mer-Cre-Mer transgenic mice. These mice were maintained in C57BL/6N background.

### O-Ring Aortic Banding (ORAB)

ORAB surgery is a new surgical technique to induce pressure-overload-induced heart failure as described previously [27]. Mice were anesthetized by inhalation of 2% isoflurane and then intubated with a 20-gauge catheter and ventilated using a volume-controlled rodent ventilator and Minivent Anesthesia Systems (Hugo Sachs Elektronik, March, Germany) with a stroke volume of 0.25 ml oxygen and isoflurane mixture and a respiratory rate of 200 breaths min^−1^. After shaving the hair on the chest, thoracotomy was performed to access the transverse aorta. The transverse aortic arch was then constricted between the innominate and left common carotid arteries with an O-ring Nitril (0.012 x 0.016 IN, Applerubber) firmly around using prolene suture (Ethicon, EH7403). The operation procedure was performed on a clean and sterile operation field.

### Generation of Adeno-Associated Viruses 9 (AAV9)

The DNA sequence encoding *Camk2D-C* isoform with an N-terminal MYC tag or firefly luciferase (Luc) was cloned into a single-stranded AAV9 vector containing the cardiac-specific troponin-T promoter and packaged into AAV9 vector. The use of cardio-trophic AAV9 vectors in addition to the use of a cardiac-specific promoter system allowed for cardiac-specific expression of *Camk2D-C*. AAV9 vectors were produced with the two-plasmid transfection method as described previously [57]. For cardiac-specific gene delivery, 8-week-old mice received the indicated AAV9 vectors through a single tail-vein injection. To allow for expression of the above-mentioned constructs, mice were kept for 4 weeks. The dosages used for the experiments were 1 × 10^12^ genomic particles.

### Lentivirus production and knockdown assay

The pLKO.1-puro constructs containing oligonucleotides for *SLC27A1* or *CD36* sequences were used. The lentiviral packaging plasmids containing pMD2.G, pRRE and pRSV-rev, along with the pLKO.1 shRNA, were transfected into HEK 293T cells using GeneJammer (Agilent). Virus-containing medium were collected 72 h after transfection and then used to infect iPS-CM. The knockdown efficiency was assessed by western blotting. The oligonucleotide sequences for the shRNA are: CD36 shRNA: GCCATAATCGACACATATAAA, and SLC27A1 shRNA: CCCTGATCTTTGGAGGAGAAA.

### Mouse echocardiography

Ventricular function was assessed by transthoracic echocardiography using a Vevo 2100 high-resolution imaging system (Visual Sonics Inc.) equipped with a MS400 transducer. Briefly, mice were shaved and measurements were performed on conscious mice. B-mode and M-mode images were recorded from the parasternal long axis and in short axis view at midpapillary muscle level. The values were analyzed with the VevoLab software.

### Reagents and adenoviruses

Isoprenaline (ISO, Sigma) was used at a concentration of 0.2 μM; 2-NBDG (2-(*N*-(7-Nitrobenz-2-oxa-1,3-diazol-4-yl)Amino)-2-Deoxyglucose) (Thermo Fischer Scientific) was used at a concentration of 300 μM; Oleic acid (Sigma) was used at a concentration of 100 μM. The NR4A1 inhibitor DIM-C-pPhOH (DIM) at a concentration of 20 μM was used as described previously [17]. The following adenoviral vectors encoding GFP-tagged *Nr4a1* (Ad.*Nr4a1*) and MYC-tagged *Camk2* (Ad.*Camk2*) constitutively active form were used to overexpress NR4A1 and *CAMK2* respectively. An adenovirus expressing GFP (Ad.GFP) alone served as the control.

### Cell Culture and transfection

iPS-CMs were maintained at 37 °C and 5% CO_2_ in RPMI medium with B27 supplements. C_2_C_12_ myoblasts and HEK293 (ATCC) cells were grown in Dulbecco’s Modified Eagle’s Medium (DMEM, Sigma [D5796]) supplemented with fetal calf serum (10%) and penicillin/streptomycin (1%). Transfection was performed with lipofectamine 2000 (Life Technologies, Invitrogen) according to the manufacturer’s instructions.

### Glucose and lipid uptake

Adult mouse ventricular myocytes (AMVMs) from cDKO or corresponding wild type (WT) mice were isolated following standard enzymatic digestion according to the Langendorff perfusion system. Cells were placed on the laminin-coated cell culture plate and maintained in fresh complete medium. AMVMs were stimulated with ISO (0.2 μM) for two hours. After stimulation in order to measure glucose uptake, the cells were washed and then incubated with 2-NBDG (300 μM) in glucose-free DMEM (Thermo Fischer Scientific) for 30 min at 37^0^C. For lipid droplet staining, AMVMs were stimulated simultaneously with ISO (0.2 μM) and Oleic acid (100 μM) for two hours. After washing, intracellular fluorescence was acquired by confocal microscopy (Leica SP8).

### Histology and immunohistochemistry (IHC)

Hearts were cut transversally for two-chamber view, washed quickly in 0.9% NaCl and fixed in 4% PBS-buffered paraformaldehyde (PFA). The hearts were then embedded in paraffin. Staining with either haematoxylin and eosin (H&E) or Masson’s trichrome was performed. For IHC, heart tissue sections were deparaffinized and then blocked in 5% BSA for 1 h at room temperature. Dilute primary antibody (1:200) anti-α-sarcomeric actinin (Sigma A7811) (anti-Perilipin2; Progen, GP40) incubation was in a humidity chamber overnight at 4 °C. Secondary antibodies (Invitrogen, conjugated with Alexa Fluor dyes) were applied for 1 h at room temperature (1:400) along with counterstaining with DAPI (1:5000). Lipid staining was performed using BODIPY (Thermo Fischer Scientific, D3922) (1:2000) for 1 h at room temperature. Photographs were acquired for histology using slide scanner Axio Scan.Z1 (Zeiss).

### RNA-Sequencing

RNA isolated from left ventricles of 14-wk-old male wild-type (WT) and cDKO mice was analyzed on a Bioanalyzer 2100 (Agilent). Total RNA was depleted from ribosomal RNA, polyA-enriched, fragmented, and paired-end sequenced at BGI (Hongkong, CN). We provide details of the computational tools and coding scripts used in the current study as an online supplement via GitHub data repository: https://github.com/mepepin/ Alignment of reads to the mm10 genome was performed using *STAR* (v2.7.10a), with differential gene expression calculated using *DESeq2* (1.32.0) within the R (4.1.2) statistical computing environment as described previously [58, 59]. We calculated differential expression from normalized read counts via Wald’s test with Bonferroni’s post hoc adjusted *P* value for each aligned and annotated gene. Statistical significance was assumed as unpaired two-tailed Bonferroni-adjusted *P* value (*FDR*) < 0.05, with a lower-stringency statistical threshold used for pathway enrichment (*P* < 0.05).

### Real-time polymerase chain reaction

Total RNA was isolated from ventricular tissue with Trizol (Invitrogen, Germany). The synthesis of cDNA was performed using the First Strand cDNA-synthesis kit following the manufactures protocol (Thermo Fischer Scientific). RT-qPCR was performed with PowerUp SYBRGreen Master Mix (Thermo Fischer Scientific) in a Lightcycler 480 II system (Roche). Each sample was run in technical duplicates. Data was analyzed using LinRegPCR version 2018.0 [60]. The list of primers shows in supplementary table.

### Western Blotting

Ventricular tissue was homogenized in a TissueLyser II (Qiagen, the Netherlands) using RIPA buffer (50 mmol/L AQ5 Tris, pH 8, 150 mmol/L NaCl, 1 mmol/L EDTA, 0.1% SDS, 0.5% Na-Deaoxycholate and 1% NP-40), with addition of protease inhibitor (S8820, MilliporeSigma) and phosphatase inhibitor (P5726, MilliporeSigma). Total protein was quantified with a colorimetric assay (Bio-Rad). On isolation, proteins from heart tissue and cultured cardiac myocytes were separated by SDS-PAGE and transferred onto polyvinylidene fluoride membranes for Western blot. For Western blots, O-GlcNAC mouse monoclonal primary antibody (Cell Signaling, CDT 110.6, 1:000), anti-Slc27a1 rabbit antibody (Bioss Antibodies, bs-10556R), anti-CD36 rabbit antibody (Thermo Fischer Scientific, PA1-16813), mouse anti-CAMK2 antibody (BD Biosciences, 611293, 1:100), mouse anti–Nur77 antibody (BD Pharmingen, 554088, 1:1000) and mouse anti-GAPDH antibody (Millipore, MAB 374, 1:10000) were used. Primary antibody incubation was followed by corresponding horseradish peroxidase–conjugated secondary anti-mouse and anti-rabbit antibodies and enhanced chemiluminescence detection.

### Luciferase Reporter Assay

To identify whether NR4A1 regulates *Slc27a1* gene expression, luciferase reporter assay was performed. Briefly, C_2_C_12_ myoblasts were co-transfected with various plasmids, including a luciferase reporter plasmid (1µg/ml, pGL3-promoter, Promega), a renilla expression vector (500 ng/ml, pRL-TK-Renilla, Promega), and a mammalian expression vector containing either GFP-tagged *Nr4a1* or GFP alone (2 µg/ml, pcDNA3). After 24 hours post-transfection, cells were lysed and the luciferase activity was measured using Fluostar Optima (BMG Labtech) and normalized for transfection efficiency to internal renilla values. The promoter of the *Slc27a1* contains NR4A1 binding site as wild type (NBRE-like motif) or mutated binding site were amplified by PCR and then cloned into the pGL3 promoter luciferase reporter vector.

### DNA-Protein Pulldown assay

To identify whether NR4A1 directly binds to the promoter of *Slc27a1*, we performed a pulldown assay using biotinylated DNA as previously described [61]. Briefly, we designed end-labeled biotinylated DNA oligonucleotides to amplify a 324 bp fragment of *Slc27a1* promoter region contains either wild type NR4A1-binding site or a mutated NR4A1-binding site. C_2_C_12_ or HEK293 Cells were transfected with plasmids expressed either GFP-tagged NR4A1 or GFP alone. For immunoprecipitation, protein lysates were diluted for equal concentrations and precleared over streptavidin-conjugated beads (Dynabeads M-280 Streptavidin, Thermo Fisher Scientific) for 2 hours at 4°C. Precleared supernatants were added to the mixture of biotinylated DNA/streptavidin-conjugated beads and incubated for 4 hours at 4°C. Afterwards, beads were washed and eluted at 95°C for 5 minutes. The eluents were immunoblotted with an antibody against NR4A1. The biotinylated *Slc27a1* primer sequence is: (Bio) GGTTCTTGTTGACCCCTGGCTG and reverse primer: ATGAGCCCTTTGCAGAGTCC.

### FDG-PET scanning

PET imaging was performed up to 60 min after intravenous injection of 4 nmol of glucose analogue 2-deoxy-2-[^18^F] fluoro-D-glucose (FDG) per mouse using an Inveon small animal PET scanner (Siemens) and a three-dimensional emission scan was performed. Images were reconstructed iteratively and analysed using standardized uptake values (SUV). Quantification was done using a region-of-interest (ROI) technique and expressed as mean SUV.

### Steady-state metabolomics profiling

Ground heart tissue was further homogenized in a Mixer Mill (Retsch) and ceramic beads at maximum frequency for 2 minutes to 4 minutes in pre-cooled racks after adding ice-cold methanol/H_2_O (4:1, v/v, 500 µL per 20 mg tissue) with internal standards. 1000 µL of homogenate was then collected in a Pyrex® glass tube and extracted by applying 120 µL 0.2 M HCl, 400 µL chloroform, 400 µL chloroform, and 400 µL H_2_O consecutively with vortexing steps in between. The extracts were spun down at 3000 g for 10 minutes, and the upper phase (water-soluble metabolites) was evaporated for 30 minutes at 35°C under nitrogen and dried in SpeedVac (Eppendorf) at 15 °C overnight. The lower phase (lipids) was evaporated to dryness at 45 °C under nitrogen. The interphase was used to determine the protein concentration with the BCA assay according to the manufacturer’s instructions. Samples were stored at -80 °C.

### LC-MS analysis of water-soluble metabolites

Water-soluble metabolites were dissolved in 100 µl 5 mM ammonium acetate (in 75% acetonitrile (v/v)) before loading to LC/MS. LC-MS analysis was performed on an Ultimate 3000 HPLC system (Thermo Fisher Scientific) coupled with a Q Exactive Plus MS (Thermo Fisher Scientific) in both ESI positive and negative mode. The analytical gradients were carried out using an Accucore 150-Amide-HILIC column (2.6 µm, 2.1 mm x 100 mm, Thermo Fisher Scientific) with solvent A (5 mM ammonium acetate in 5% acetonitrile) and solvent B (5 mM ammonium acetate in 95% acetonitrile). 3 µl sample was applied to the Amide-HILIC column at 30°C, and the analytical gradient lasted 20 minutes. During this time, 98% of solvent B was applied for 1 minute, followed by a linear decrease to 40% within 5 minutes, which was maintained for 13 minutes before returning to 98% within 1 minute and remaining in starting conditions for 5 minutes for equilibration. The flow rate was maintained at 350 µL/min. The eluents were analyzed with MS in ESI positive/negative mode with ddMS2. The full scan at 70k resolution (69-1000 m/z scan range, 1e6 AGC-Target, 50 ms maximum Injection Time (maxIT)) was followed by a ddMS2 at 17.5k resolution (1e5 AGC target, 50 ms maxIT, 1 loop count, 0.1 s to 10 s apex trigger, 2e3 minimum AGC target, 20 s dynamic exclusion). The HESI source parameters were set as 30 sheath gas flow rate, 10 auxiliary gas flow rate, 0 sweep gas flow rate, spray voltage: 3.6 kV in positive mode, 2.5 kV in negative mode, 320 °C capillary temperature, and the heater temperature of auxiliary gas was 120 °C. The annotation of the metabolites was performed using the EI-Maven software (Elucidata, https://www.elucidata.io/el-maven) with an offset of ± 15ppm.

### LC-MS/MS analysis of lipids

Lipids were dissolved in 100 µl of isopropylalcohol (iPrOH) before loading. The analytical gradients were carried out using an Accucore C8 column (2.6 µm, 2.1 mm x 50 mm, Thermo Fisher Scientific) with solvent A (acetonitrile/H_2_O/formic acid (10/89.9/0.1, v/v/v)) and solvent B (acetonitrile/iPrOH/H_2_O/formic acid (45/45/9.9/0.1, v/v/v/v)). 3 µl sample were applied to the C8 column at 40°C, and the analytical gradient lasted for 35 minutes. During this time, 20% of solvent B was applied for 2 minutes, followed by a linear increase to 99.5% within 5 minutes and maintained for 27 minutes before returning to 20% in 1 minute and appended with a 5-minute equilibration step. The flow rate was maintained at 350 µL/min. The full scan and ddMS2 parameters were the same as the analysis of the water-soluble metabolites, except the scan range were adjusted to 200-1600 m/z. The HESI source parameters were also adapted with a 3 sweep gas flow rate and a 3.2 kV spray voltage in positive and 3.0 kV in negative mode. Peaks corresponding to the calculated lipid masses (± 5 ppm) were integrated using El-Maven software (Elucidata, https://www.elucidata.io/el-maven).

### Statistical Analyses

All studies were performed on at least 3 independent occasions. Results are reported as mean ± SEM. Statistical analysis was performed with GraphPad Prism Software Package version 6.0 (GraphPad Inc) The comparison of different groups was carried out with a 2-tailed unpaired Student *t* test or 2-way ANOVA, followed by a post hoc Bonferroni or Tukey test. A value of *P*<0.05 was considered statistically significant.

**Supplementary Table S1.**
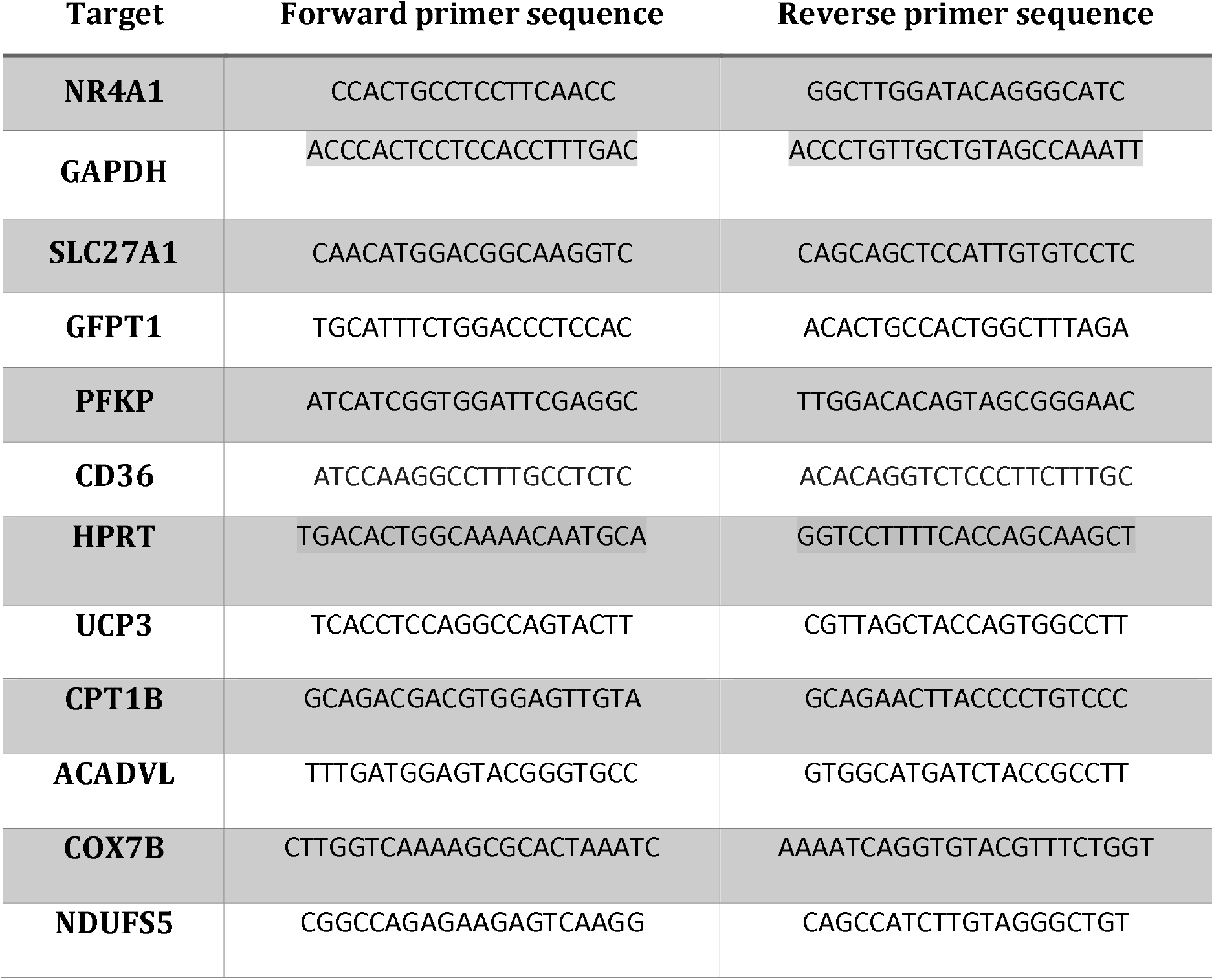
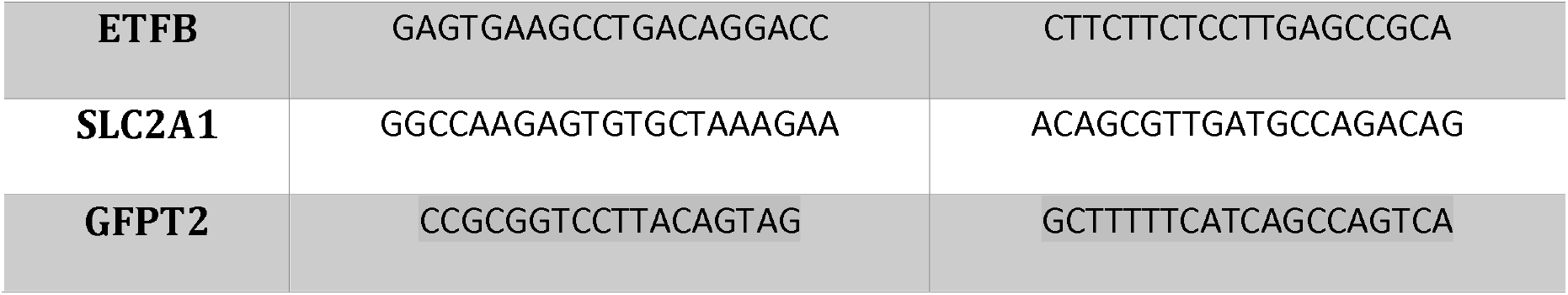
Primer Sequences in human species

**Supplementary Table S2.**
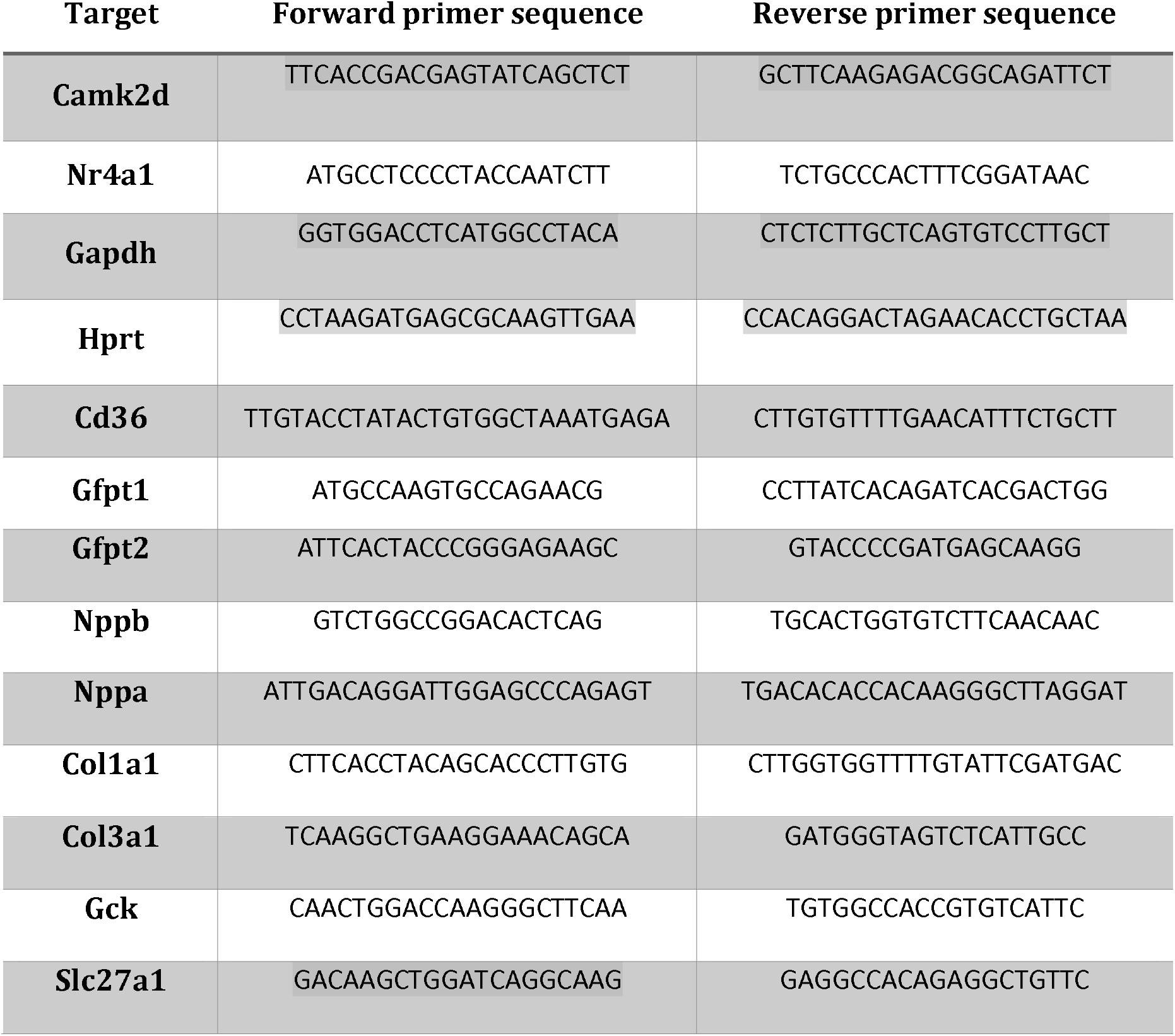
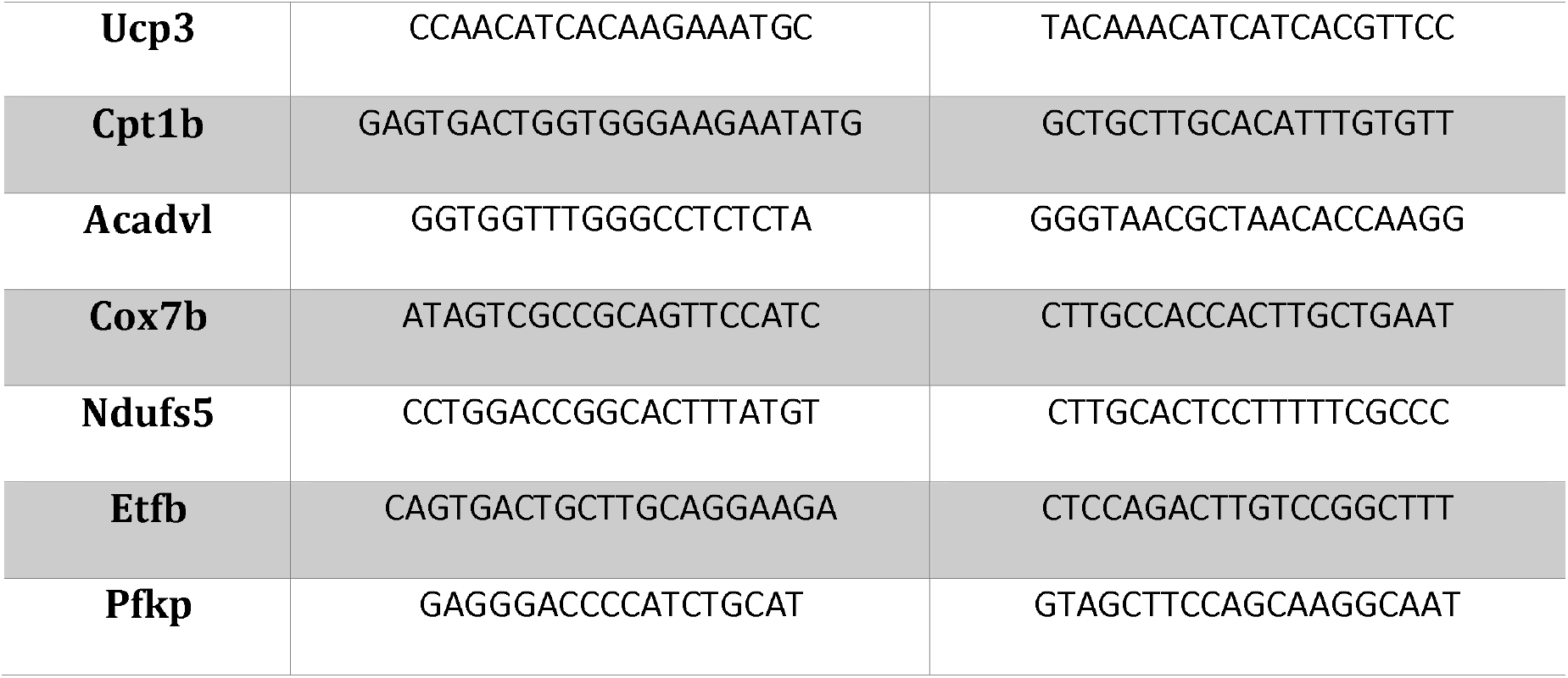
Primer Sequences mouse species

